# The *Escherichia coli* chromosome moves to the replisome

**DOI:** 10.1101/2023.07.12.548795

**Authors:** Konrad Gras, David Fange, Johan Elf

**Affiliations:** Department of Cell and Molecular Biology, Science for Life Laboratory, Uppsala University, Husargatan 3, Uppsala 75124, Sweden

## Abstract

The replisome, a large multi-subunit assembly, carries out the process of chromosome replication, connecting the unwrapping of the parental DNA with the creation of new daughter strands. In *Escherichia coli*, it is debated whether the two replisomes move independently along the two chromosome arms during replication or if they remain in close proximity, with the DNA being pulled toward the replisomes as replication progresses. Here, we use fluorescence microscopy to determine the location and diffusivity of the replisome and various chromosomal loci throughout the cell cycle of the model organism *E. coli*. We find that (*i*) the two replisomes are confined to a region of 250 nm and 120 nm along the cell long and short axis respectively, and the chromosomal loci move through this region sequentially based on distance from the origin of replication. (*ii*) When a locus is being replicated, its diffusivity slows down. (*iii*) There is no indication that replication initiation occurs close to the cell membrane as has been proposed in a few previous studies. In conclusion, our data supports a model with DNA moving towards stationary replisomes at replication.

## Introduction

The organization of the chromosome in bacteria is influenced by various essential processes, including DNA replication, segregation, and gene expression. In *Escherichia coli*, the chromosome is compacted along the length of the cell with coiled loops extending from an axial core (Mäkelä and Sherratt 2020; Kavenoff and Ryder 1976), forming a bottle-brush-like structure (Le *et al*., 2013). While being highly coiled to fit into the bacterial cell, chromosomal DNA has to be replicated and segregated before cell division to ensure propagation into progenies. The protein complex responsible for replicating DNA, the replisome (O’Donnell, 2006; Kurth and O’Donnell, 2009), initiates replication at the origin of replication, *oriC*, and replicates the circular chromosome sequence bidirectionally towards the terminus region, *ter*. Contrary to the sequential aspects of chromosome replication in *E. coli*, the spatial aspects of this process are still under debate. Various studies have presented differing models for if, how, and where the replisome acts on the chromosome structure. One model describes the replisome as relatively stationary during replication, with DNA moving through the replisome (Hiraga *et al*., 2000; Molina and Skarstad, 2004; Adachi *et al*., 2005; Mangiameli *et al*., 2017). An alternative model instead describes the chromosomal DNA as stationary, with the replisome moving along the DNA (Kongsuwan *et al*., 2002; Bates and Kleckner, 2005; Reyes-Lamothe *et al*., 2008; Japaridze *et al*., 2020). Stationary replisomes have also been observed in other bacterial organisms, such as *Bacillus subtilis* (Lemon and Grossman, 1998; Berkmen and Grossman, 2006; Mangiameli *et al*., 2017) and *Caulobacter crescentus* (Jensen, Wang and Shapiro, 2001). Combinations of the two models have been proposed, although with contradictory suggestions of how cells combine the different replication modes (Espeli, Mercier and Boccard, 2008; Chen *et al*., 2023). The experimental evidence that supports the respective models in *E. coli* has been previously discussed (Mangiameli *et al*., 2018), suggesting that differences in the data analysis could have given rise to the contradicting models. Models involving mobile replisomes propose that the replication forks initiate in the middle of the cell and then separate by traveling bidirectionally along the DNA until they meet at *ter*. In contrast, models that describe the replisome as stationary suggest that DNA is pulled through the replisomes, which, for certain models (Sawitzke and Austin, 2001), have been speculated to be anchored or coupled to each other.

Knowing where replication occurs is crucial for understanding the growth-dependent organization of the *E. coli* chromosome. Here we revisit the question of how the localization of a locus is affected by replication. To do so, we used high-throughput fluorescence microscopy of live *E. coli* cells to simultaneously study both replication and chromosome localization. Initially, we focused our efforts on the origin of replication, *oriC*, which by physical necessity colocalizes with the replisome at replication initiation. We measured the diffusivity of *oriC* over the cell cycle and observed a decrease in the diffusivity of the origin region at replication initiation. A similar behavior was observed for loci further away from *oriC*, with the diffusivity decrease occurring as the loci were pulled into the more stationary replisome. Our results strongly suggest that the chromosome moves in response to being replicated while the replisomes stay in a confined intracellular region during replication.

## Results

### An *oriC*-proximal chromosome locus shows more dynamics over the cell cycle as compared to the replisome

To investigate if the chromosome or the replisome, or both, relocate during the replication process, we constructed a strain with fluorescent labels of different colors on the replisome and an *oriC*-proximal chromosome locus. Initially, we focused on *oriC* since initiation occurs within a narrow range of cell sizes (Knöppel *et al*., 2023), making interpretation more straightforward. We engineered an *E. coli* MG1655 strain with DnaN translationally fused with mCherry (Moolman *et al*., 2014). We also introduced a YFP-based fluorescent repressor-operator system (FROS) (Lau *et al*., 2004) 34 kb from *oriC*. The FROS-based locus labeling was performed with an array of 12 *malO* operators and constitutive expression of the gene encoding MalI-SYFP2. We grew the strain in a mother-machine-type microfluidic chip (Baltekin *et al*., 2017), where we imaged the bacteria every minute over several hours. Cell outlines were estimated using phase-contrast microscopy and assembled into tracks of cell lineages based on the time-lapsed acquisitions. Fluorescence images of the replisome and *oriC* labels were acquired back-to-back with two 25 ms laser excitation pulses of different colors, <1 ms apart. The split emission channels were projected onto different parts of a single camera chip.

Using back-to-back imaging of our origin and replisome labels, we aimed to investigate how *oriCs* localize over the cell cycle and, in particular, at replication initiation. To visualize where the replisome and the *oriC*-proximal loci are positioned within the cells, each replisome and locus fluorescent focus was placed in a cell internal coordinate system defined by the cell outline captured using phase-contrast. The area enclosed by the cell outline is hereafter referred to as cell area. The cells were binned based on cell area and for each bin, we generated bi-variate distributions of the long- and short-axis intracellular positions of replisomes and *oriC*-proximal loci (Figure 1).

**Figure 1.**
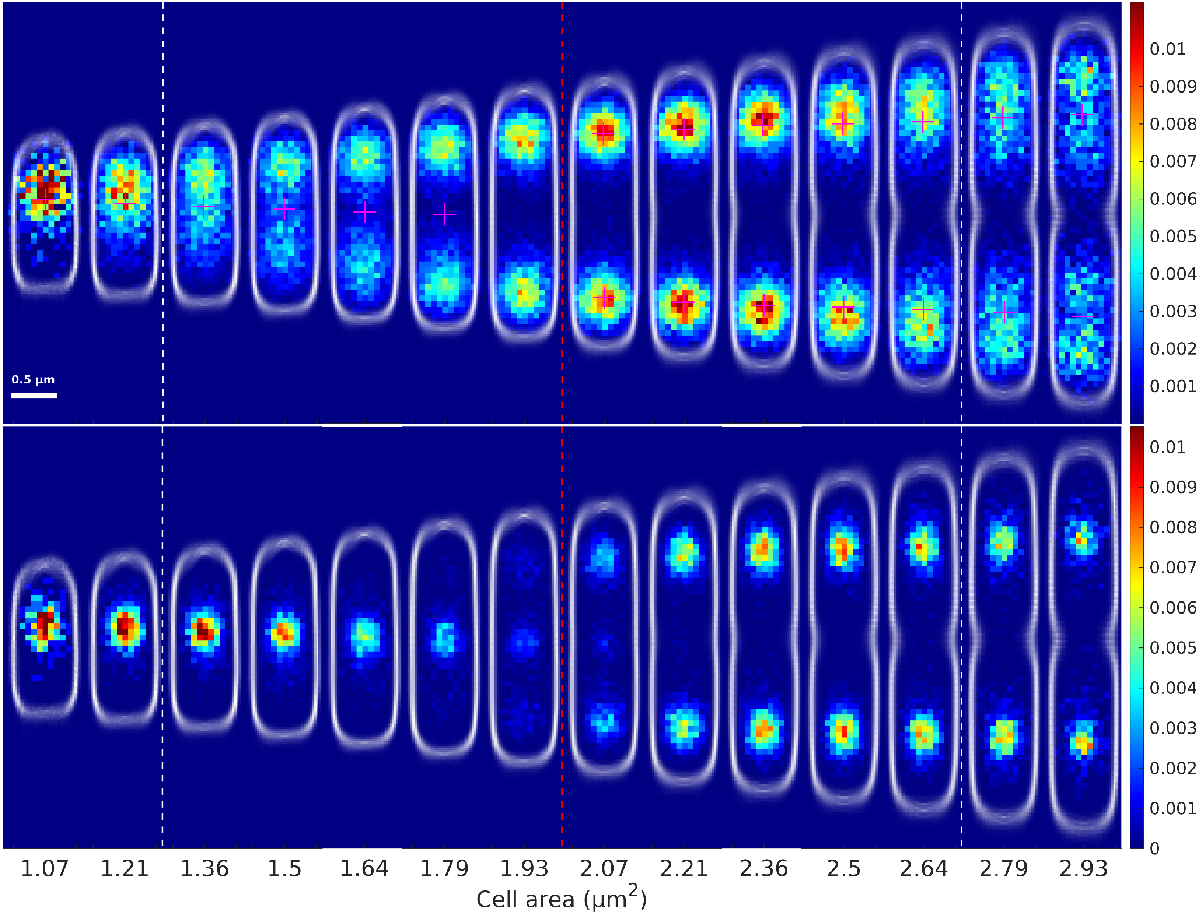
*OriC* and replisome localization distributions over the cell cycle. Two-dimensional histograms of fluorescent foci positions along the long and short cell axis from an *E. coli* strain that simultaneously carries a chromosomal locus label (*malO*/MalI system with an array of operator sequences introduced 34 kb from *oriC*, top) and a replisome label (mCherry-DnaN, bottom). Color in each histogram heat-map indicates the number of foci per cell and histogram-voxel, where all the two-dimensional histograms use the same voxel-size and voxel positions. The distributions of cell-outlines detected in phase-contrast are shown in white. The old pole is always pointing up. White dashed lines indicate average birth and division sizes. Red dashed lines indicate average replication initiation size. The average birth, division and replication initiation sizes have been adjusted to fit in between the two-dimensional histograms. Magenta plus signs correspond to the mean of the replisome distributions from the same strain based on 2D Gaussian fitting of the distribution. Figures are generated from foci in 86930 cells tracked from division to subsequent division.

At a cell area of ∼2 μm^2^, two replisome distributions spawn around the 1/4 and ¾ long-axis positions of the cell (Figure 1, bottom). We interpret these spawns as the initiation of two new rounds of replication on the origins already in place at these positions (Wallden et al, 2016). Thus, each of the replisome distributions corresponds to a pair of replication forks replicating the chromosomal DNA sequence bidirectionally away from *oriC* (Reyes-Lamothe *et al*., 2008; Wallden *et al*., 2016). As expected, the average of the localization distribution of the *oriC*-proximal chromosome loci colocalizes with the replisome distribution in the size span where initiation occurs (Figure 1, top). During one generation, the replicated *oriCs* must segregate into the daughter cells. We find that the segregation of oriC occurs for most of the time between two initiations The replisome distributions, on the other hand, stay localized at a similar average distance from the middle of the cell throughout the replication process. Note that after cell division, the old cell middle now equals the new pole. Before termination, the replisomes stay between the two segregating *oriC* distributions (Figure 1). Although there are two replication forks in each of the two replisome distributions, the width of the distributions remains relatively constant (Figure S1), indicating that the two replisomes moving bidirectionally on the chromosome stay close together throughout the replication process (Mangiameli *et al*., 2017).

The replisome localization distributions in Figure 1 could arise either as a result of stationary replisomes with cell-to-cell variability in replisome localization, or from replisomes moving around, either individually or as pairs of bidirectionally moving replication forks. To discriminate between these scenarios, we tracked the replisomes in individual cells for 20 minutes following the initiation event and recorded how the root mean squared displacement (RMSD) changed in time (Figure 2A). The RMSDs plateau at ∼0.25 μm in the cell long-axis direction and ∼0.12 μm in the cell short-axis direction. The plateau values for both the short and long axis of the cell are similar to the expected average distance between two random positions in the replisome localization distributions (Figure 2B), suggesting that the major contributor to the width of the replisome distributions shown in Figure 1 is replisome movements and not cell-to-cell variation.

**Figure 2.**
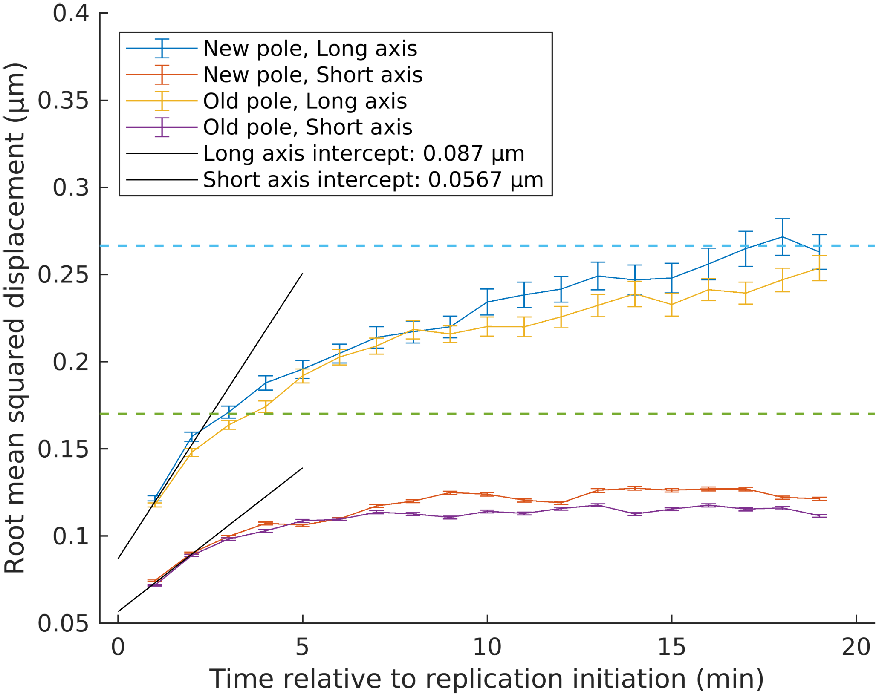
Replisome RMSD following the replication initiation event. RMSD of the replisome over 20 minutes relative to the replication initiation event. Blue and green dashed lines indicate √2 * standard deviation from 2D Gaussian fits of the replisome localization distributions along the long and short axis respectively. Black lines correspond to linear fits of the first two RMSD values for the long and short axes. Error bars correspond to standard error of the mean.

Taken together, we find that the localization distribution of the replisome varies less compared to the *oriC*-proximal locus, both in terms of its average position and width of the distribution. These localization patterns are indicative of stationary replisomes and moving chromosomal loci. However, they are inconclusive in terms of movements immediately after the initiation event, since small changes in the localization of the *oriC*-proximal locus are easily lost in the diffusive movements of both the locus and the replisomes.

### Replication of a locus coincides with a decrease in its diffusivity

To further characterize the movement of the *oriC*-proximal locus and the replisome, we measured their diffusivity. This was done by acquiring *oriC* and replisome images back-to-back, repeating the process after 1 second and estimating the displacement of the fluorescent foci over the 1 second interval. The distributions of frame-to-frame displacement at different cell sizes for both *oriC* and replisomes are shown in Figure 3A and 3B. We found that the diffusivity of the *oriC*-proximal loci decreases at the cell size window where initiation occurs, while the replisome diffusivity stays constant and low over the cell cycle. Measurements of two-dimensional (2D) Euclidean distances between the replisome and *oriC*-proximal loci for different cell areas (Figure 3C) also show that the displacement minimum coincides with the minimal replisome-*oriC* distance.

**Figure 3.**
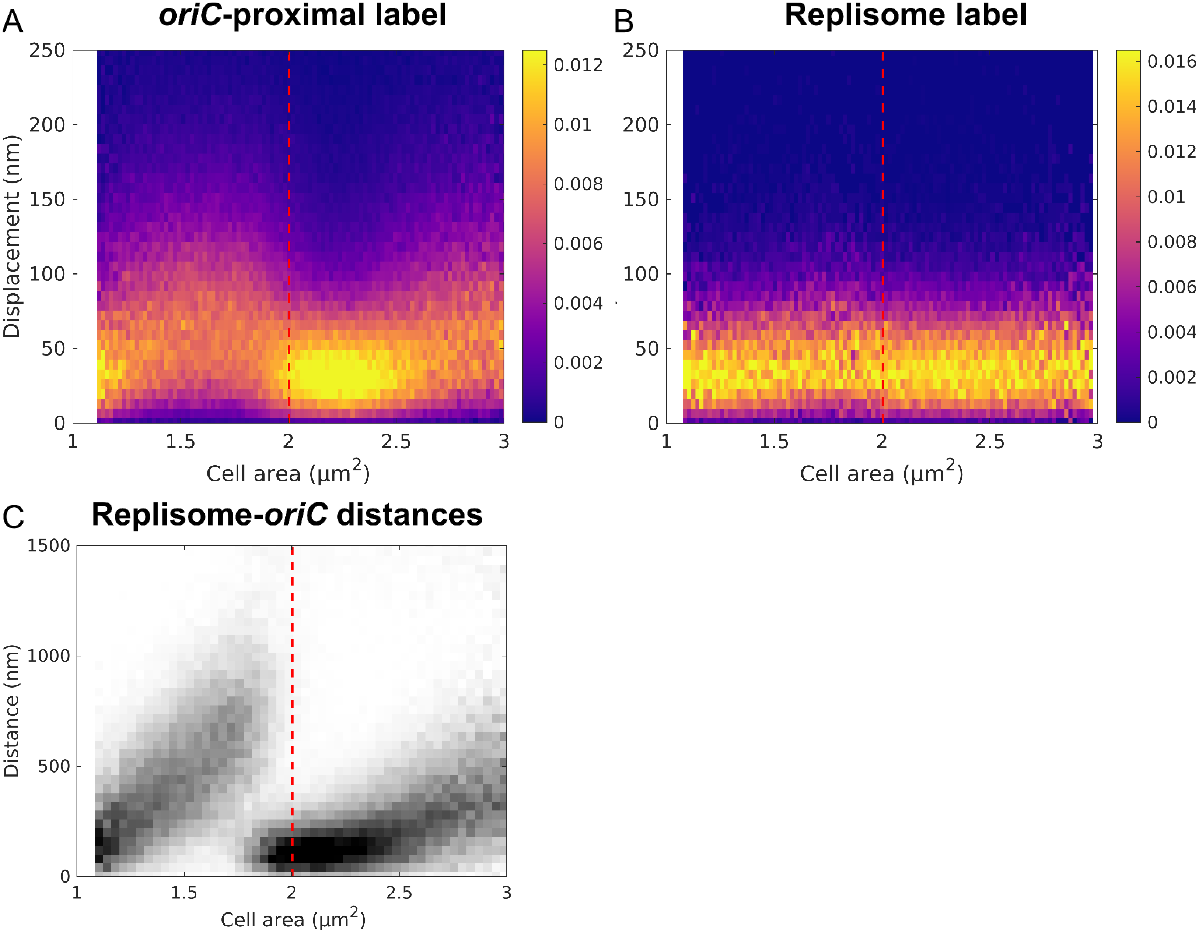
*OriC* and replisome diffusivity at different cell areas. **(A & B)** Frame-to-frame displacement of tracked fluorescent foci between two frames acquired 1 s apart. Tracking was performed using the u-track (Jaqaman *et al*., 2008) algorithm. Displacements at different cell sizes estimated for **(A)** *oriC*-proximal chromosome loci and **(B)** replisomes. **(C)** 2D distances between the *oriC-*proximal locus and replisomes for different cell areas. Red dashed lines indicate average replication initiation size.

If the decrease in replisome-*oriC* displacements and the replisome-*oriC* distance minimum are due to the replication fork passing the locus, we should see a corresponding pattern for other chromosomal loci at well-defined times after initiation. To test this, we measured displacements at 1-s time intervals for 11 strains with the YFP-based FROS labels at different distances from *oriC*. In all 11 strains, we observed displacement minima similar to that of the *oriC*-proximal labeled locus, but with the minima occurring at different cell sizes (Figure 4). The minima of loci displacements occur at the same cell area as the minima in replisome-locus distance, which, in turn, are consistent with a model where initiation occurs at 2.05 μm^2^, replication takes 45 minutes to complete, progressing at a constant rate, and exponentially growing cells with a doubling time of 50 minutes (see Materials & Methods for details). These results show that as loci are replicated, their diffusivity decreases to that of the replisome (Figure 4).

**Figure 4.**
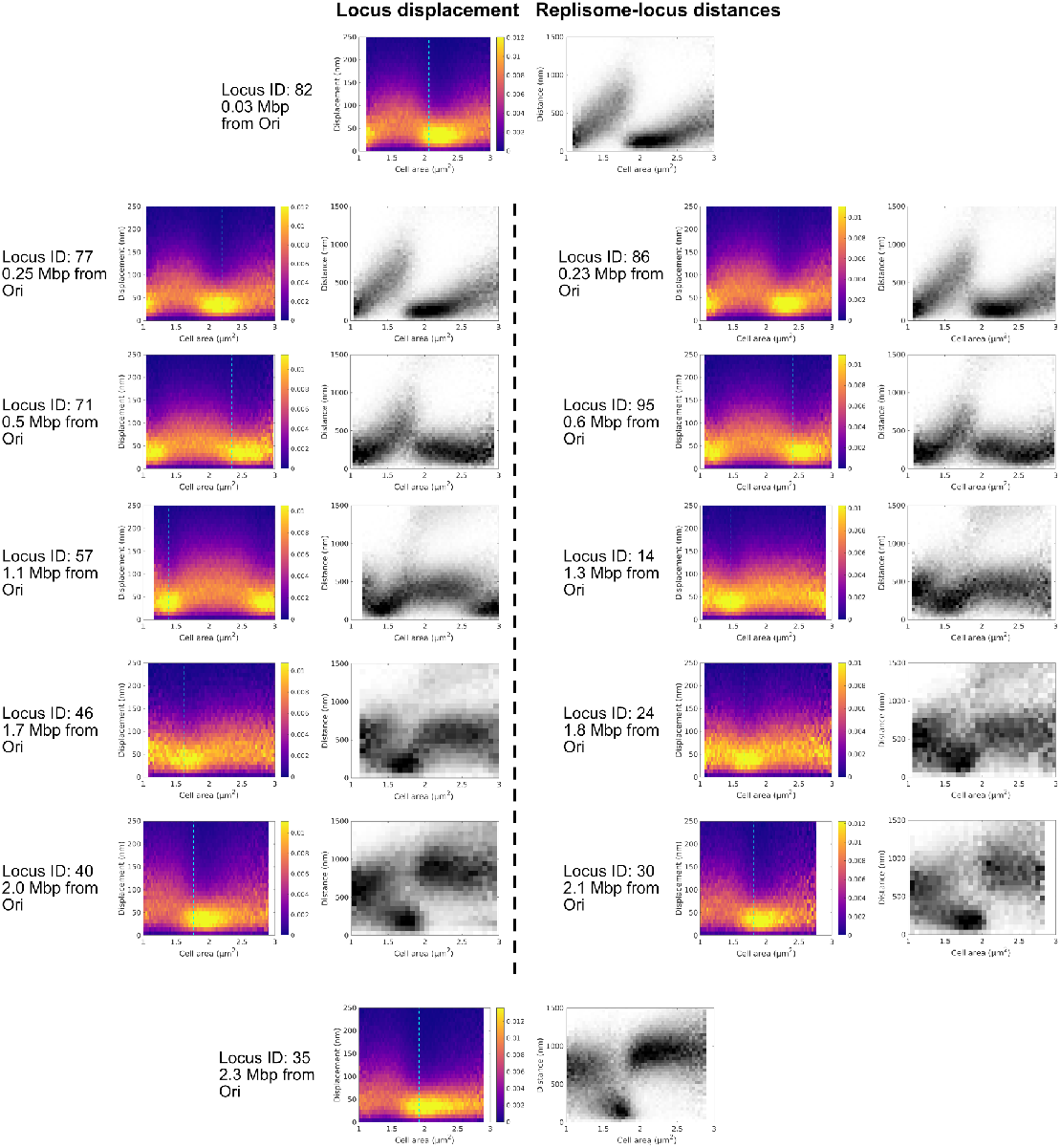
Locus displacement minima coincide with replisome-locus colocalization. As Figure 3A and 3C but for chromosome locus labels at different chromosomal positions relative to *oriC*. Presented data for Locus ID: 82 is the same as in Figure 3A and 3C. Cyan lines indicate expected average cell size of replication based on a model where initiation occurs at 2.05 μm^2^, replication progresses at a constant rate and takes 45 minutes to complete, and exponentially growing cells with a doubling time of 50 minutes.

### Replication induces transient spatial repositioning of chromosomal loci

Using the 12 strains with FROS labels in different positions, we also quantified the diffusivity at different spatial positions inside each cell (Figure 5). We note that minima occur when the localization distributions of each locus and the replisome overlap, in agreement with our distance measurements. For the *oriC*-proximal label, we find an overall expected behavior, with a decrease in displacement at the cell size interval where initiation occurs, but without a clear intracellular position-dependent diffusivity. However, for chromosomal loci which are 1.7 and 1.8 Mbp away from *oriC* on the left and right chromosomal arms respectively, an interesting behavior emerges. At cell sizes of ∼1.6 μm^2^, the loci relocate from the old pole into the region of the replisome, in which the locus diffusivity becomes slower. Shortly after replication, these loci are segregated into their new positions and remain there until the next replication round. As the loci segregate, their diffusivity returns to the pre-replication level. These observations show that the chromosome moves as a result of the replication process and not vice versa.

**Figure 5.**
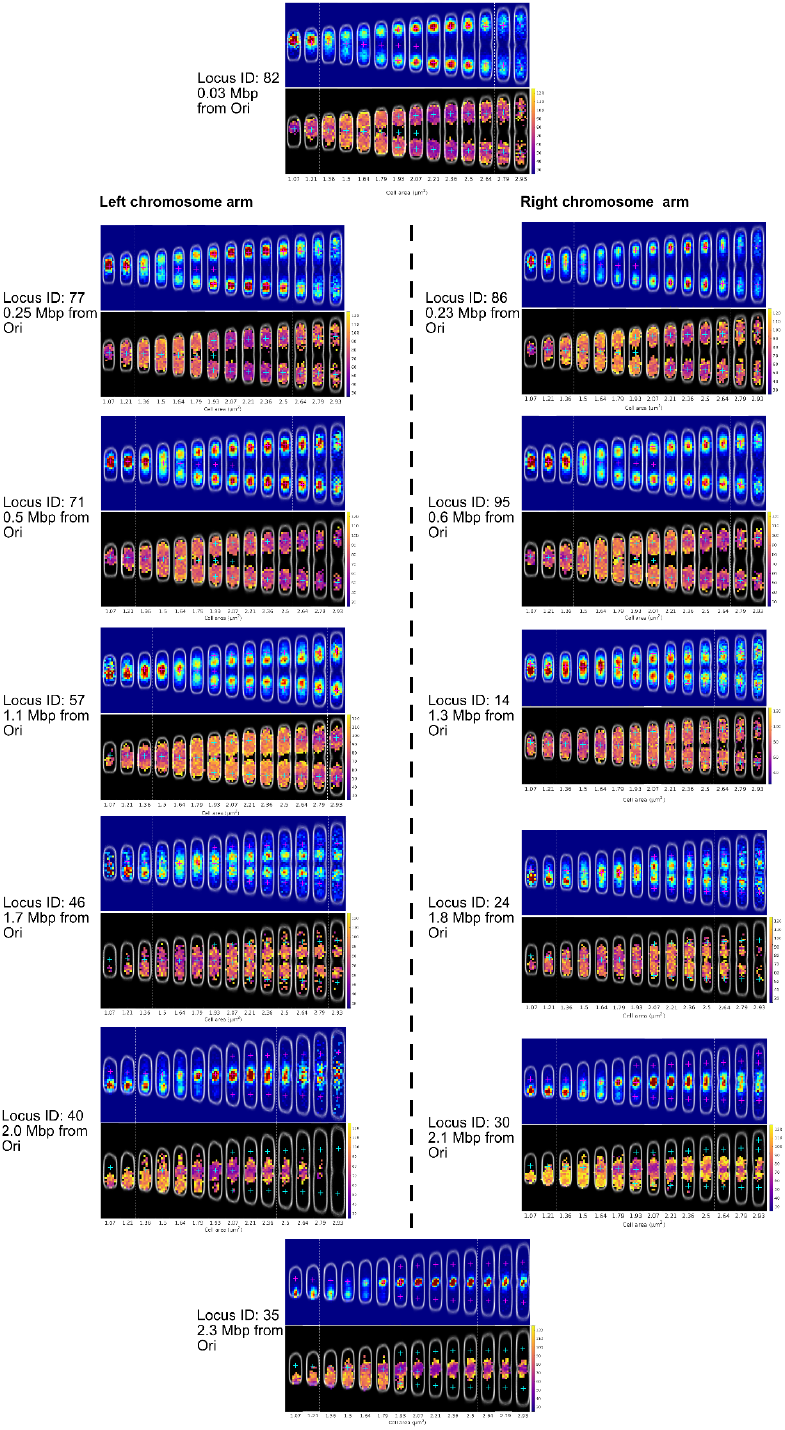
Locus diffusivity at different spatial positions. Two-dimensional histograms of fluorescent foci positions and displacements along the long and short axes of the cell. Histograms based on focus positions as in Figure 1, but for chromosome locus labels at different chromosomal positions relative to *oriC*. Two-dimensional histograms of focus displacements over 1 s at different spatial positions in the cell are based on the same positions as the focus localizations. The displacement histogram heat-map color indicates the mean displacement at each spatial position based on at least 5 displacements measured at that position. Magenta and cyan plus signs correspond to the mean of a 2D Gaussian fit to the replisome distributions from the same strain.

### Initiation of replication does not occur at the cell membrane

A number of studies have implicated components of the cell membrane as being involved in regulating DNA replication. For example, acidic phospholipids present in the cell membrane have been suggested to be involved in regulating chromosome replication initiation (Saxena *et al*., 2013). The binding of these phospholipids to *oriC* also blocks *oriC* binding of the DnaA protein (Crooke, Castuma and Kornberg, 1992), which, in its ATP form, is required for replication initiation (Sekimizu, Bramhill and Kornberg, 1987). Following initiation, DnaA is converted to its ADP form, which by itself is not sufficient to initiate replication (Crooke, Castuma and Kornberg, 1992). Hence, DnaA needs to be converted into its ATP form to trigger initiation again. The acidic phospholipids have, *in vitro*, been shown to facilitate this conversion for *oriC*-bound DnaA-ADP (Sekimizu, Bramhill and Kornberg, 1987; Yung, Crooke and Kornberg, 1990; Castuma, Crooke and Kornberg, 1993). Hemimethylated *oriC*, which temporarily exists in the wake of the replication fork, has also been shown to bind to various components of the *E. coli* cell membrane (Hendrickson *et al*., 1982); (Kusano *et al*., no date; Chakraborti *et al*., 1992). The interaction has been suggested to depend on SeqA, which binds hemimethylated DNA after it has been replicated. Furthermore, the interaction between *oriC* and the cell membrane might impact the ability to initiate replication (Landoulsi *et al*., 1990). The membrane affinity has also been used to formulate a model where replication initiation is triggered based on the detachment of *oriC* from the membrane (Norris, 1990). Taken together, these observations indicate that *oriC* should be located near the membrane, especially close to replication initiation.

Contrary to the membrane involvement suggested by previous studies, we did not observe localization distributions reminiscent of membrane binding at any point in the cell cycle (Figure 1). The lack of membrane-proximal localization applied to both the replisome and *oriC*-proximal labels. The distribution of replisomes shown in Figure 1 is even excluded from the membrane regions as identified by phase contrast (Figure 1). To highlight what localization distribution we should expect from a membrane-bound protein, we imaged LacY-PAmCherry and localized it as a function of cell size (Figure 6A). A comparison of the short-axis distribution of the replisome and LacY for cell areas close to the average initiation size shows that the overlap between these two distributions is very small (Figure 6B). To ensure that we were not missing any transient membrane-binding events, we also investigated localization distributions of the replisome and *oriC*-proximal labels as a function of time relative to replication initiation (Figure 6C). However, we could not observe any membrane-proximal localization patterns in this case either. Taken together, our results show that replication initiation does not occur at the cell membrane.

**Figure 6.**
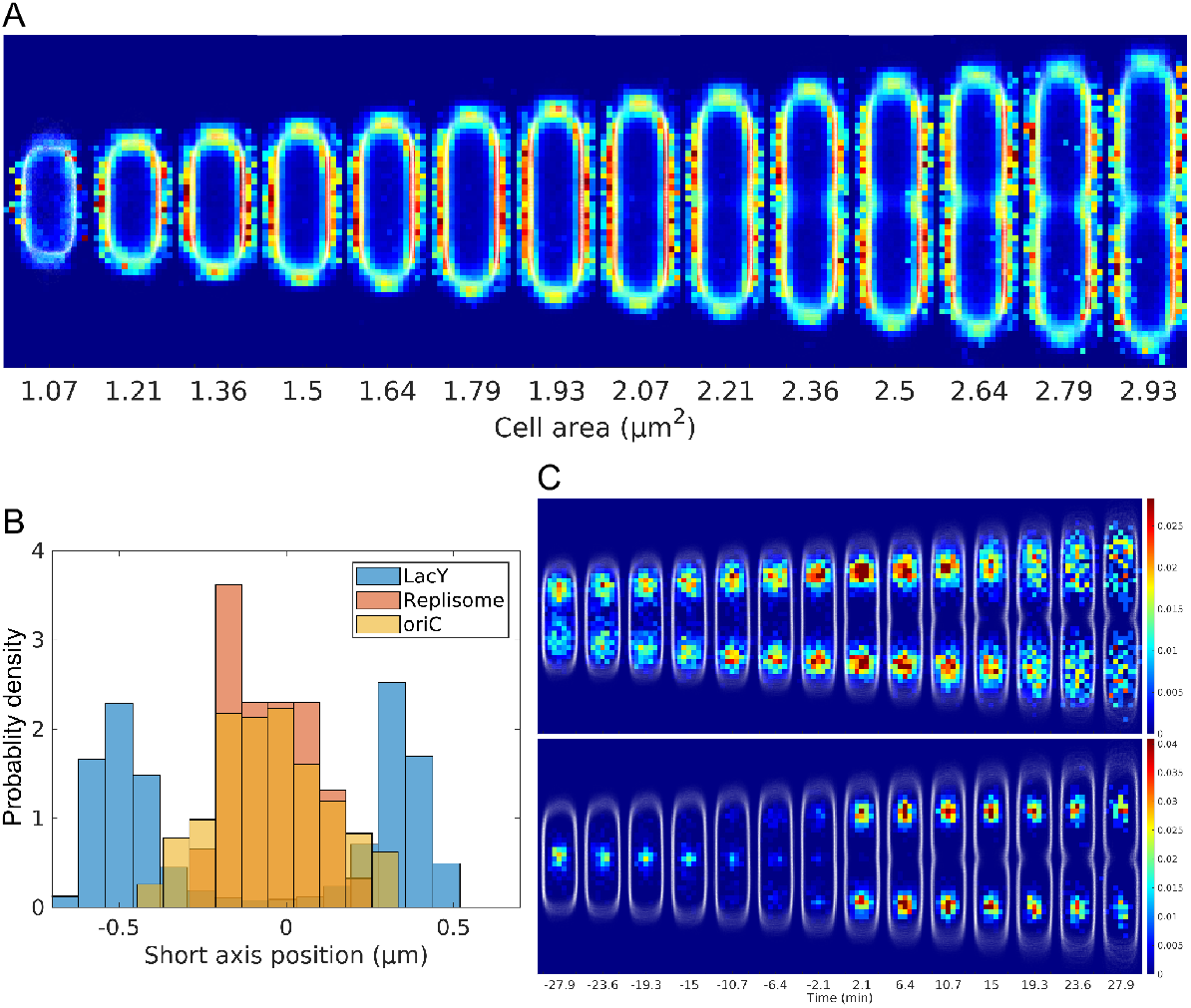
Comparing the localization distributions of LacY, *oriC* and the replisome. **(A)** Localization distributions as in Figure 1 but for LacY-PAmCherry. **(B)** Histograms of short-axis localization coordinates of LacY-PAmCherry, the replisome and *oriC*-proximal labels around the ¼ position along the cell at ∼2 μm^2^. **(C)** Localization distributions as in Figure 1, but sorted based on time relative to replication initiation.

## Discussion

We show striking evidence that chromosomal loci make large intracellular movements both directly before and directly after replication while the replisomes stay relatively stationary. This behavior is most clear for the loci 1.7 Mbp and 1.8 Mbp from *oriC*. These loci translocate, on average, more than half of the cell length during a period of 30% of the average generation time (Figure 5, Locus ID: 24 & 46). During the same period, the average movement of the replisomes is 5% of a cell length. The *ter*-proximal loci exhibit similar pre-replicative movement, but show no post-replicative segregation since their replication occurs close to the intracellular space at which *ter* resides for a majority of the cell cycle. Rapid relocalization of *ter* has been reported previously (Espeli, Mercier and Boccard, 2008), and it has been suggested that this process is a direct consequence of replication (Espéli *et al*., 2012). The notion that the loci move while the replisomes are stationary is in line with observations of replisomes in (Hiraga *et al*., 2000; Molina and Skarstad, 2004; Adachi *et al*., 2005; Mangiameli *et al*., 2017), but possibly contradictory to the conclusions from direct concurrent observations of replisomes and chromosome loci in the same cells made in (Reyes-Lamothe *et al*., 2008) and recently by (Japaridze *et al*., 2020).

Based on replisome localization distributions (Figure 1) and replisome MSD (Figure 2), we find that replisomes stay spatially bounded within intracellular regions with a standard deviation width of 0.12 μm in the cell short axis direction and 0.25 μm in the cell long axis direction. The size of the replisome region also remains relatively unchanged throughout the replication cycle (Figure S1), in line with previous observations (Wallden *et al*., 2016; Mangiameli *et al*., 2018; Knöppel *et al*., 2023). The two replisomes can, within the region, temporarily be detected as separate (Mangiameli *et al*., 2017; Knöppel *et al*., 2023). Although it has been shown that it is possible for the replisomes to move apart for extended periods of time (Reyes-Lamothe *et al*., 2008), most of the time they are detected as one fluorescent focus (Mangiameli *et al*., 2017). The transient presence of multiple foci together with the observations made by (Japaridze *et al*., 2020) that the replisomes move apart in cells that have an altered cell geometry indicate that bi-directionally moving replisomes do not form stable multi-replisome complexes while replicating.

Our displacement measurements (Figure 3) show that the chromosomal loci have higher diffusivity compared to the replisome. This is in contrast to results reported in (Reyes-Lamothe *et al*., 2008), where the opposite relation was described. A potential explanation for this discrepancy could be differences in labeling strategy, as our FROS labels are based on shorter arrays (12 repeats of binding sites as compared to 240). We also find that locus diffusivity depends on the intracellular position, which is in line with (Javer *et al*., 2013). Our data suggest that the replication process is the underlying reason for this dependency for a majority of the chromosomal loci.

Among the loci that we investigated, the *ter*-proximal locus exhibits a specifically distinct diffusivity pattern over the cell cycle (Figure 4 & 5, Locus ID: 35, 47 kb from *dif*). The decrease in diffusivity of other loci is temporary, i.e. occurring only during replication, while *ter* remains relatively immobile at mid-cell after replication until cell division. The observation of slower diffusion for ter-proximal loci is also in line with previous observations (Javer *et al*., 2013). As the replisome and *ter* colocalize, replication terminates, which results in catenated copies of *ter* (Steiner *et al*., 1999; Li, Sergueev and Austin, 2002). The combined activity of XerCD, FtsK, and Topoisomerase IV at the *dif* site mediate unlinking of the *ter* regions (Hojgaard *et al*., 1999; Steiner *et al*., 1999). The catenation of this region could contribute to its low diffusivity and narrow localization distribution, compared to other loci. Notably, FtsK has been implicated in chromosome localization (Liu, Draper and Donachie, 1998) and segregation of *ter* (Stouf, Meile and Cornet, 2013). Additionally, it has been shown that the *ter*-specific DNA-binding protein MatP helps to localize the terminus region at mid-cell through an interaction with the divisome proteins ZapB, ZapA, and FtsZ (Espéli *et al*., 2012; Männik *et al*., 2016). The interactions with a number of cell division components unique to the *ter* region likely contribute to its distinctive displacement patterns.

It has previously been suggested that stationary bacterial replisomes could facilitate the segregation of newly replicated loci by extruding them towards the cell poles (Lemon and Grossman, 1998, 2001; Sawitzke and Austin, 2001). Replication as the main driver for chromosome segregation has previously been discounted by the observation that loci do not segregate immediately after replication (Bates and Kleckner, 2005). Contrary to this, our results show that for most loci investigated, the localization distributions do appear to split following a decrease in displacement (Figure 5), which we interpret as locus segregation following its replication. Notable exceptions are loci close to *oriC* and *ter*. However, determining when a locus is segregated based on localization of fluorescent foci alone is challenging, as two copies of a locus can be colocalized in a diffraction-limited focus. Thus, the replication of a locus may not appear to be immediately followed by its segregation due to the locus labeling, and not as a result of delayed segregation. Therefore, a replication-driven model of chromosome segregation cannot be ruled out.

A conversion of DnaA-ADP → DnaA-ATP facilitated by acidic phospholipids could provide the missing piece of the puzzle for how the replication initiation potential is increased prior to initiation (Knöppel *et al*., 2023). Computational modeling indicates that to provide stable initiation events at well-defined cell sizes, the phospholipid-assisted conversion of ADP to ATP on DnaA should occur when DnaA is not bound to the chromosome (Berger and Wolde, 2022). In contrast, *in vitro* data suggests that the process should involve the chromosome. If *oriC* was membrane localized at initiation, this would have provided a clear indication of the involvement and function of phospholipids in initiation. However, we see no evidence of membrane localization of the origin region at any point in the cell cycle (Figures 1 & 6). Any transient membrane interactions would have to occur on a very short time scale. Given our observations (Figure 6), a direct link between DnaA-ADP at *oriC* and the phospholipids in the membrane seems unlikely.

In this study, we have shown that various regions on the *E. coli* chromosome exhibit higher diffusivity than the replisomes. Replication takes place within a restricted region into which loci move when they are replicated. During the replication process, locus diffusivity decreases and becomes similar to the replisome diffusivity. We observe several cases of loci moving towards the replisome to be replicated and subsequently segregated. Our results also show that replication initiation does not occur at the cell membrane. To conclude, these observations support a model with a stationary and confined replisome region through which the chromosome moves to be replicated.

## Materials & Methods

### Strains and growth conditions

All constructed strains were derivatives of the strain *E. coli* MG1655 *rph*+ *ΔmalI::frt ΔintC::P59-malI-SYFP2-frt ΔgtrA::SpR frt*-*mCherry*-*dnaN*. For the construction of these strains chromosomal modifications were performed using λ Red recombineering (Datsenko and Wanner, 2000) and generalized transduction with phage P1 *vir* (Thomason, Costantino and Court, 2007). All experiments were performed in M9 minimal medium supplemented with 0.06× Pluronic F-108 (Sigma-Aldrich 542342) 0.4% succinate and 1× RPMI 1640 amino acid solution (Sigma) at 30 °C. Strains were inoculated one day before each experiment in culture tubes with growth medium from frozen stock cultures stored at −80 °C. Cells were grown overnight at 30 °C in a shaking incubator (200 rpm). On the day of the experiment, the cells were diluted 1:100 in growth medium and grown for 3 to 4 h before being loaded in the microfluidic chip.

A mother machine-type PDMS chip with open-ended channels was used for all microfluidic experiments. The width and height of these channels was 1000 nm. Medium flow and loading of cells into the chip was performed by supplying pressure to the chip ports with a microfluidic flow controller built in-house (AnduinFlow). The imaging of the chip was performed in a H201-ENCLOSURE hood that enclosed the microscope stage, with the hood being connected to a H201-T-UNIT-BL temperature controller (OKOlab).

### Microscopy

All imaging was performed with a Ti-E (Nikon) microscope equipped with a 100X immersion oil objective (Nikon, CFI Apo Lambda, NA 1.45) that was set up for phase-contrast and widefield epi-fluorescence microscopy. Phase-contrast and fluorescence images were acquired of cells growing in a microfluidic chip at 30 °C in M9 minimal medium supplemented with 0.4% succinate and 1x RPMI amino acids solution (Sigma). The imaging was performed over 10 h, unless otherwise noted. Phase-contrast and fluorescence images were acquired once per position on the microfluidic device with a 1 min^-1^ imaging frequency. Each imaged position included 20 traps with approximately 200-300 cells in total per position.

For fluorescence image acquisition, the sample was irradiated with a 580 nm laser (VFL, MBP Communications) for mCherry excitation followed by a 515 nm laser (Fandango 150, Cobolt) for SYFP2 excitation. Both lasers were set to a 25 ms exposure time and 30 W/cm^2^ power density at the image plane. The laser light was shuttered using an AOTFnC together with MDPS (AA Opto-Electronic) and reflected onto the sample using a FF444/521/608-Di01 (Semrock) triple-band dichroic mirror. Fluorescence images were acquired using a Kinetix sCMOS camera (Teledyne Photometrics). Imaging of the two fluorescence channels was performed back-to-back by two function generators (Tektronix) that triggered the lasers based on the camera acquisition. Emitted fluorescence was transmitted through a BrightLine FF580-FDi02-T3 (Semrock) dichroic beamsplitter to separate the fluorescence for different channels. The separated fluorescence was then filtered through BrightLine FF01-505/119-25 (Semrock) and BrightLine FF02-641/75-25 (Semrock) filters and focused on two different areas of the sCMOS camera.

For the displacement measurements the fluorescence imaging also involved an additional image acquisition in the two fluorescence channels at a given time interval after the first acquisition. The double acquisitions with a time interval were performed by using the SMART Streaming feature of the sCMOS camera, which allows for sequential acquisition of images with different exposure times. One of the acquisitions was used only to introduce the time interval between the two frames used for the displacement measurements.

Phase-contrast images were acquired with a 50 ms exposure time using a DMK 38UX304 camera (The Imaging Source). The light source used for phase-contrast was a 480 nm LED and a TLED+ (Sutter Instruments). The transmitted light was passed through the same FF444/521/608-Di01 (Semrock) triple-band dichroic mirror as the fluorescence and reflected onto the camera with a Di02-R514 (Semrock) dichroic mirror.

Imaging of LacY-PAmCherry used the same microscopy and microfluidics setup as for the strains with replisome and locus labels. For fluorescence image acquisition the sample was irradiated with stroboscopic illumination of a 580 nm laser (VFL, MBP Communications) with 30 ms pulses for fluorescence excitation and a 405 nm laser (Cobolt) with 7.5 ms pulses for fluorophore activation. The illumination was triggered using two function generators (Tektronix) based on the camera acquisition.

### Image analysis

The image analysis was performed using an automated image analysis pipeline that is primarily written in MATLAB R2022a (Mathworks) and previously described in (Camsund *et al*., 2020). Cell segmentation of phase-contrast images was performed using a nested Unet neural network (Ronneberger, Fischer and Brox, 2015). Cell tracking was performed using the Baxter algorithm (Magnusson *et al*., 2015) and tracking of fluorescent foci was performed using the u-track algorithm (Jaqaman *et al*., 2008). Fluorescent foci of SYFP2 and mCherry were detected using a wavelet-based detection algorithm (Olivo-Marin, 2002) and the detected coordinates were then refined using a maximum likelihood-based localization algorithm (Lindén *et al*., 2017). The localization was performed with fitting of a two-dimensional (2D) Gaussian function to the fluorescent signal. To transform the localized focus coordinates between the cameras used to acquire fluorescence and phase-contrast images, landmark-based registration was performed of 500 nm beads (TetraSpeck, Thermo Fisher) that were visible on both cameras.

The estimation of replisome-*oriC* distances involved pairing of localized mCherry-DnaN and MalI-SYFP2 foci from the same cell. landmark-based registration was performed by imaging 100 nm fluorescent beads (TetraSpeck, Thermo Fisher) in two different fluorescence channels simultaneously on two different areas of the camera chip. The pairing between foci from two different fluorescence channels was performed based on this registration. Distances were estimated based on foci that were the closest to each other and each focus was only paired with one other focus from the other fluorescence channel. The pairing of foci coordinates was performed using MATLAB’s matchpairs() function.

Predicated replication sizes for the fluorescently labeled loci were calculated assuming exponential growth of the cells and using an average replication initiation size of 2.05 μm^2^, an average generation time of 50 min and an average C-period of 45 min. The replication size A(t) is then estimated by *A*(α) = *A*_*Init*_ * *exp*(μ * *C* * α), where A_*init*_ is the average replication initiation size, μ is the average growth rate, C is the average C-period and ∝ is the genomic distance of a given locus from *oriC* divided by half of the length of the chromosome.

## Acknowledgements

We thank Irmeli Barkefors for critical reading of the manuscript. Nynke Dekker provided us with strains containing the mCherry-DnaN construct. Elias Amselem built the two color setup. This study was made possible by grants from the ERC (advanced grant no. 885360), the Swedish Research Council (grant nos. 2016-06213 and 2018-03958), the Knut and Alice Wallenberg Foundation (grant nos. 2016.0077, 2017.0291, and 2019.0439). The computations and data management were enabled by resources provided by the Swedish National Infrastructure for Computing at UPPMAX, partially funded by the Swedish Research Council through grant agreement no. 2018-05973.

## Author contributions

D.F., and J.E. conceived the study; K.G., J.E. and D.F. designed experiments; K.G. constructed most strains, performed the experiments and analyzed the data; K.G. and D.F. wrote most of the analysis code; K.G., J.E. and D.F. interpreted results; K.G., J.E. and D.F. wrote the paper.

---

**Figure S1.**
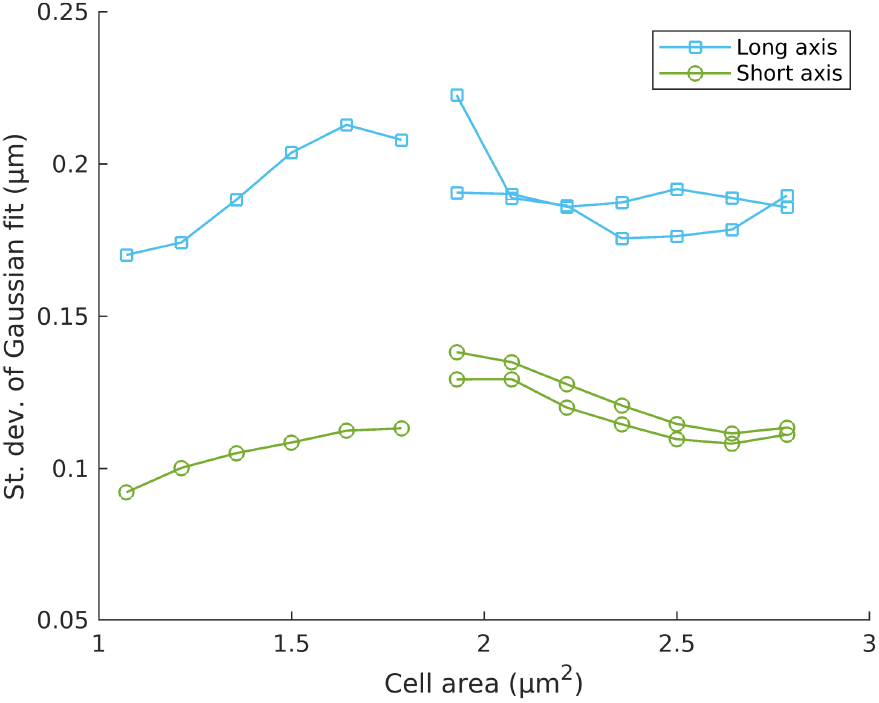
Standard deviations of replisome localization distributions. Standard deviations based on 2D Gaussian fits of replisome localization distributions in Figure 1 as a function of cell size.

## References

1. Adachi, S. et al. (2005) ‘Localization of replication forks in wild-type and mukB mutant cells of Escherichia coli’, Molecular genetics and genomics: MGG, 274(3), pp. 264–271.

2. Baltekin, Ö. et al. (2017) ‘Antibiotic susceptibility testing in less than 30 min using direct single-cell imaging’, Proceedings of the National Academy of Sciences of the United States of America, 114(34), pp. 9170–9175.

3. Bates, D. and Kleckner, N. (2005) ‘Chromosome and replisome dynamics in E. coli: loss of sister cohesion triggers global chromosome movement and mediates chromosome segregation’, Cell, 121(6), pp. 899–911.

4. Berger, M. and Wolde, P.R.T. (2022) ‘Robust replication initiation from coupled homeostatic mechanisms’, Nature communications, 13(1), p. 6556.

5. Berkmen, M.B. and Grossman, A.D. (2006) ‘Spatial and temporal organization of the Bacillus subtilis replication cycle’, Molecular microbiology, 62(1), pp. 57–71.

6. Camsund, D. et al. (2020) ‘Time-resolved imaging-based CRISPRi screening’, Nature methods, 17(1), pp. 86–92.

7. Castuma, C.E., Crooke, E. and Kornberg, A. (1993) ‘Fluid membranes with acidic domains activate DnaA, the initiator protein of replication in Escherichia coli’, The Journal of biological chemistry, 268(33), pp. 24665–24668.

8. Chakraborti, A. et al. (1992) ‘Characterization of the Escherichia coli membrane domain responsible for binding oriC DNA’, Journal of bacteriology, 174(22), pp. 7202–7206.

9. Chen, P.J. et al. (2023) ‘Interdependent progression of bidirectional sister replisomes in E. coli’, eLife, 12. Available at: https://doi.org/10.7554/eLife.82241.

10. Crooke, E., Castuma, C.E. and Kornberg, A. (1992) ‘The chromosome origin of Escherichia coli stabilizes DnaA protein during rejuvenation by phospholipids’, The Journal of biological chemistry, 267(24), pp. 16779–16782.

11. Datsenko, K.A. and Wanner, B.L. (2000) ‘One-step inactivation of chromosomal genes in Escherichia coli K-12 using PCR products’, Proceedings of the National Academy of Sciences, 97(12), pp. 6640–6645.

12. Espéli, O. et al. (2012) ‘A MatP–divisome interaction coordinates chromosome segregation with cell division in E. coli’, The EMBO journal, 31(14), pp. 3198–3211.

13. Espeli, O., Mercier, R. and Boccard, F. (2008) ‘DNA dynamics vary according to macrodomain topography in the E. coli chromosome’, Molecular microbiology, 68(6), pp. 1418–1427.

14. Hendrickson, W.G. et al. (1982) ‘Binding of the origin of replication of Escherichia coli to the outer membrane’, Cell, 30(3), pp. 915–923.

15. Hiraga, S. et al. (2000) ‘Bidirectional migration of SeqA-bound hemimethylated DNA clusters and pairing of oriC copies in Escherichia coli’, Genes to cells: devoted to molecular & cellular mechanisms, 5(5), pp. 327–341.

16. Hojgaard, A. et al. (1999) ‘Norfloxacin-induced DNA cleavage occurs at the dif resolvase locus in Escherichia coli and is the result of interaction with topoisomerase IV’, Molecular microbiology, 33(5), pp. 1027–1036.

17. Japaridze, A. et al. (2020) ‘Direct observation of independently moving replisomes in Escherichia coli’, Nature communications, 11(1), p. 3109.

18. Jaqaman, K. et al. (2008) ‘Robust single-particle tracking in live-cell time-lapse sequences’, Nature methods, 5(8), pp. 695–702.

19. Javer, A. et al. (2013) ‘Short-time movement of E. coli chromosomal loci depends on coordinate and subcellular localization’, Nature communications, 4, p. 3003.

20. Jensen, R.B., Wang, S.C. and Shapiro, L. (2001) ‘A moving DNA replication factory in Caulobacter crescentus’, The EMBO journal, 20(17), pp. 4952–4963.

21. Kavenoff, R. and Ryder, O.A. (1976) ‘Electron microscopy of membrane-associated folded chromosomes of Escherichia coli’, Chromosoma, 55(1), pp. 13–25.

22. Knöppel, A. et al. (2023) ‘Regulatory elements coordinating initiation of chromosome replication to the Escherichia coli cell cycle’, Proceedings of the National Academy of Sciences of the United States of America, 120(22), p. e2213795120.

23. Kongsuwan, K. et al. (2002) ‘Cellular localisation of the clamp protein during DNA replication’, FEMS microbiology letters, 216(2), pp. 255–262.

24. Kurth, I. and O’Donnell, M. (2009) ‘Replisome Dynamics during Chromosome Duplication’, EcoSal Plus, 3(2). Available at: https://doi.org/10.1128/ecosalplus.4.4.2.

25. Kusano, T. et al. (no date) ‘Direct evidence for specific binding of the replicative origin of the Escherichia coli chromosome to the membrane’, Journal of bacteriology [Preprint].

26. Landoulsi, A. et al. (1990) ‘The E. coli cell surface specifically prevents the initiation of DNA replication at oriC on heminethylated DNA templates’, Cell, 63(5), pp. 1053–1060.

27. Lau, I.F. et al. (2004) ‘Spatial and temporal organization of replicating Escherichia coli chromosomes: Escherichia coli chromosome dynamics’, Molecular microbiology, 49(3), pp. 731–743.

28. Lemon, K.P. and Grossman, A.D. (1998) ‘Localization of Bacterial DNA Polymerase: Evidence for a Factory Model of Replication’, Science, 282(5393), pp. 1516–1519.

29. Lemon, K.P. and Grossman, A.D. (2001) ‘The extrusion-capture model for chromosome partitioning in bacteria’, Genes & development, 15(16), pp. 2031–2041.

30. Le, T.B.K. et al. (2013) ‘High-resolution mapping of the spatial organization of a bacterial chromosome’, Science, 342(6159), pp. 731–734.

31. Lindén, M. et al. (2017) ‘Pointwise error estimates in localization microscopy’, Nature communications, 8(1), p. 15115.

32. Liu, G., Draper, G.C. and Donachie, W.D. (1998) ‘FtsK is a bifunctional protein involved in cell division and chromosome localization in Escherichia coli’, Molecular microbiology, 29(3), pp. 893–903.

33. Li, Y., Sergueev, K. and Austin, S. (2002) ‘The segregation of the Escherichia coli origin and terminus of replication’, Molecular microbiology, 46(4), pp. 985–996.

34. Magnusson, K.E.G. et al. (2015) ‘Global linking of cell tracks using the Viterbi algorithm’, IEEE transactions on medical imaging, 34(4), pp. 911–929.

35. Mäkelä, J. and Sherratt, D.J. (2020) ‘Organization of the Escherichia coli Chromosome by a MukBEF Axial Core’, Molecular cell, 78(2), pp. 250–260.e5.

36. Mangiameli, S.M. et al. (2017) ‘The Replisomes Remain Spatially Proximal throughout the Cell Cycle in Bacteria’, PLoS genetics, 13(1), p. e1006582.

37. Mangiameli, S.M. et al. (2018) ‘The bacterial replisome has factory-like localization’, Current genetics, 64(5), pp. 1029–1036.

38. Männik, J. et al. (2016) ‘The role of MatP, ZapA and ZapB in chromosomal organization and dynamics in Escherichia coli’, Nucleic acids research, 44(3), pp. 1216–1226.

39. Molina, F. and Skarstad, K. (2004) ‘Replication fork and SeqA focus distributions in Escherichia coli suggest a replication hyperstructure dependent on nucleotide metabolism’, Molecular microbiology, 52(6), pp. 1597–1612.

40. Moolman, M.C. et al. (2014) ‘Slow unloading leads to DNA-bound β2-sliding clamp accumulation in live Escherichia coli cells’, Nature communications, 5, p. 5820.

41. Norris, V. (1990) ‘DNA replication in Escherichia coli is initiated by membrane detachment of oriC’, Journal of molecular biology, 215(1), pp. 67–71.

42. O’Donnell, M. (2006) ‘Replisome architecture and dynamics in Escherichia coli’, The Journal of biological chemistry, 281(16), pp. 10653–10656.

43. Olivo-Marin, J.-C. (2002) ‘Extraction of spots in biological images using multiscale products’, Pattern recognition, 35(9), pp. 1989–1996.

44. Reyes-Lamothe, R. et al. (2008) ‘Independent Positioning and Action of Escherichia coli Replisomes in Live Cells’, Cell, 133(1), pp. 90–102.

45. Ronneberger, O., Fischer, P. and Brox, T. (2015) ‘U-Net: Convolutional Networks for Biomedical Image Segmentation’. arXiv. Available at: http://arxiv.org/abs/1505.04597 x(Accessed: 5 April 2023).

46. Sawitzke, J. and Austin, S. (2001) ‘An analysis of the factory model for chromosome replication and segregation in bacteria’, Molecular microbiology, 40(4), pp. 786–794.

47. Saxena, R. et al. (2013) ‘Crosstalk between DnaA protein, the initiator of Escherichia coli chromosomal replication, and acidic phospholipids present in bacterial membranes’, International journal of molecular sciences, 14(4), pp. 8517–8537.

48. Sekimizu, K., Bramhill, D. and Kornberg, A. (1987) ‘ATP activates dnaA protein in initiating replication of plasmids bearing the origin of the E. coli chromosome’, Cell, 50(2), pp. 259–265.

49. Steiner, W. et al. (1999) ‘The cytoplasmic domain of FtsK protein is required for resolution of chromosome dimers’, Molecular microbiology, 31(2), pp. 579–583.

50. Stouf, M., Meile, J.-C. and Cornet, F. (2013) ‘FtsK actively segregates sister chromosomes in Escherichia coli’, Proceedings of the National Academy of Sciences of the United States of America, 110(27), pp. 11157–11162.

51. Thomason, L.C., Costantino, N. and Court, D.L. (2007) ‘E. coli genome manipulation by P1 transduction’, Current protocols in molecular biology / edited by Frederick M. Ausubel … [et al.], Chapter 1, pp. 1.17.1–1.17.8.

52. Wallden, M. et al. (2016) ‘The Synchronization of Replication and Division Cycles in Individual E. coli Cells’, Cell, 166(3), pp. 729–739.

53. Yung, B.Y., Crooke, E. and Kornberg, A. (1990) ‘Fate of the DnaA initiator protein in replication at the origin of the Escherichia coli chromosome in vitro’, The Journal of biological chemistry, 265(3), pp. 1282–1285.

